# ^13^C tracing in synaptosomes reveals that SGLT2 inhibition with dapagliflozin prevents metabolic deficits in the 5X-FAD model of Alzheimer’s Disease

**DOI:** 10.1101/2025.04.30.651373

**Authors:** Marisa Mekkittikul, Cansheng Zhu, Bezawit T. Danna, Xiaohong Zuo, Amy Rios, Krista Yang, Miranda Villanueva, Devanshi Agarwal, Daniel Castro, Xuelin Gu, Alexander R. Strofs, Orian S. Shirihai, Linsey Stiles, Sally A. Frautschy, Jan Oscarsson, Russell L. Esterline, Gregory M. Cole, Ajit S. Divakaruni

## Abstract

Metabolic dysfunction is linked to several forms of age-related neurodegeneration including Alzheimer’s Disease (AD), and targeting brain energy metabolism is an increasingly attractive mode of therapeutic intervention. However, commonly used *in vitro* methods to identify specific metabolic pathways of interest in preclinical models of neurodegenerative disease have considerable limitations. They are prone to subselection of sample material, unable to identify cell type-specific effects, or cannot identify metabolic defects upstream of mitochondria. Here we address these challenges by validating a method for stable isotope tracing with isolated synaptic nerve terminals, or ‘synaptosomes’. We further applied this approach to study glucose metabolism in synaptosomes isolated from the 5X-FAD mouse model of AD treated with the antidiabetic sodium-glucose linked transporter-2 (SGLT-2) inhibitor Dapagliflozin. Treatment with Dapagliflozin preserved steady-state levels of synaptosomal metabolites and enrichment from labeled glucose into citrate that was reduced in the 5X-FAD model. These changes correlated with trends towards improved spatial working memory but not amyloid burden. The results highlight the utility of stable isotope tracing in synaptosomes to identify precise sites of metabolic dysfunction and mechanisms of action for metabolic drug candidates in preclinical models of neurodegeneration.

## INTRODUCTION

Clinical and epidemiological data strongly link cognitive impairment with aberrant glucose metabolism (1–4). Glucose hypometabolism, as diagnosed by ^18^FDG-PET scanning, can be observed years and sometimes decades prior to clinically measurable cognitive impairment (5, 6). Epidemiological data also consistently place Type 2 Diabetes at or near the top of lifestyle-related risk factors associated with Alzheimer’s Disease (AD) (7, 8). Pathology links these two diseases as well: the AD brain is insulin resistant and diabetic pancreatic islets exhibit amyloidosis (9). As such, methods to identify regulatory metabolic nodes and processes affected by disease progression in pre-clinical models are essential for both understanding the etiology of disease as well as assessing the therapeutic potential of metabolic intervention.

However, functional analysis to pinpoint specific metabolic deficits in pre-clinical neurodegenerative disease models remains underdeveloped, particularly for methods that can identify neuronal-specific effects. Non-invasive imaging methods such as PET imaging or NMR spectroscopy are highly valuable for identifying global metabolic alterations *in vivo*, but have limited utility in localizing precise mechanisms of metabolic dysfunction (10, 11). Conversely, functional analysis in mitochondria isolated from rodent brains is a powerful experimental technique for precisely identifying mechanisms of action (12, 13). However, the technique has several drawbacks including the inability to identify metabolic defects that lie upstream of mitochondria, the lack of physiologically relevant assay conditions, and the heterogeneity of the sample material which contains both astrocytic and neuronal mitochondria (14, 15).

Of course, metabolic analysis of primary neuronal cultures balances the shortcomings of imaging approaches and reductionist biochemistry. These preparations are often done with rodent embryos so the isolated cell bodies can regrow processes and synaptic junctions after weeks in culture (16–18). Unfortunately, similarly reproducible and reliable preparation of neurons from aged preclinical models of neurodegeneration can be prohibitively challenging (19), and it is unclear to what extent the surviving cell bodies isolated from adult animals are prone to subselection. Similar to cell-based analysis, isolated synaptic nerve terminals from adult rodent brains (‘synaptoneurosomes’ or ‘synaptosomes’) captures activity from both cytoplasmic and mitochondrial metabolism, and previous work has measured glucose-driven respiration in isolated synaptosomes isolated from neurodegenerative disease models (20, 21). As with all cellular respirometry studies, though, the technique cannot pinpoint defects in specific metabolic enzymes or identify altered flux through specific pathways that do not change bulk oxygen consumption rates (22).

We therefore developed methods to study glucose metabolism in synaptosomes using ^13^C stable isotope tracing. The technique allows for the direct assessment of glucose metabolism – from uptake to oxidation – in material isolated acutely and directly from the brains of adult rodents. Moreover, measurements of metabolite abundance and ^13^C enrichment with GC/MS allow for the precise localization of metabolic disruptions, particularly when these assays are coupled to complementary techniques such as respirometry in plasma membrane-permeabilized synaptosomes (23–25). In principle, the use of acutely isolated synaptosomes as opposed to primary neuronal cultures could also capture beneficial neuronal effects from physiological or pharmacologic interventions that systemically correct peripheral metabolism in pre-clinical animal models.

We applied this approach to study the systemic effects of sodium-glucose linked transporter-2 (SGLT-2) inhibition on subsequently isolated synaptosomal glucose metabolism in the 5X-FAD mouse model of Alzheimer’s Disease. The 5X-FAD model reproduces the hypometabolism of glucose observed in early human disease (26, 27), and is characterized by dysfunctional mitochondrial bioenergetics and insulin signaling (28, 29). Given the links between AD and insulin resistance, antidiabetic drugs such as SGLT-2 inhibitors are currently under clinical evaluation for neuroprotective effects (30, 31). These are of particular interest given the remarkable cardioprotective effects of SGLT-2 inhibition in patients and the potential to stimulate autophagic clearance of protein aggregates (32, 33).

Here we show metabolomics and stable isotope tracing in isolated synaptosomes can identify precise sites of metabolic dysfunction in preclinical models of neurodegenerative disease. Validation experiments show that synaptosomes exhibit appropriate signal linearity with sample material, mass isotopologue distributions, and responses to known metabolic effectors. In studying bioenergetics and glucose metabolism in 5X-FAD animals treated for 8 weeks with the SGLT-2 inhibitor Dapagliflozin, we observed that SGLT-2 inhibition resulted in a trend towards restoring spatial working memory deficits in 5X-FAD female animals. This was not associated with a rescue of amyloid burden, but GC/MS analysis revealed Dapagliflozin treatment increased steady-state levels of TCA cycle metabolites and enrichment from glucose into citrate in synaptosomes from 5X-FAD females. Altogether, the work highlights the feasibility of stable isotope tracing in isolated synaptosomes to identify metabolic pathways in preclinical disease models responsive to drugs and drug candidates.

## RESULTS

### Method Development and Validation

We first conducted a series of validation experiments to determine whether metabolite levels and stable isotope tracing data could be obtained from isolated synaptic nerve terminals (**Fig. 1A**). Indeed, metabolites from synaptosomes could be reliably detected within a linear range when loading between 75-300μg of total synaptosomal protein (**Fig. 1B**). We therefore incubated 150μg of synaptosomal protein with uniformly labeled ^13^C_6_-glucose to then determine if we could observe expected patterns of enrichment from glucose into intermediary metabolism (**Fig. 1C**). Previous bioenergetic analysis of isolated synaptosomes showed a very low ATP demand in the absence of exogenous stimuli and a relatively slow glycolytic rate that limited provision of pyruvate to mitochondria (21). We therefore supplemented our assay medium with 0.5 mM pyruvate (to overcome glycolytic rate limitations) and 1μM veratridine (a voltage-gated Na^+^-channel opener to increase the ATP demand) along with 10 mM ^13^C_6_-glucose to assess metabolite enrichment. The mass isotopologue distribution in synaptosomal metabolites showed expected patterns after a 90 min incubation. For example, lactate had a robust M+3 isotopologue, and citrate, malate, and glutamate had pronounced labeling in the evenly numbered isotopologues (e.g. M+2, M+4, associated with incorporation of the two-carbon acetyl CoA) and marginal labeling for M+1 (**Fig. 1D**). Enrichment was also observed in the M+3 and M+5 isotopologues for citrate, malate, and glutamate, highlighting that neurons are capable of glucose anaplerosis (41).

**Figure 1.**
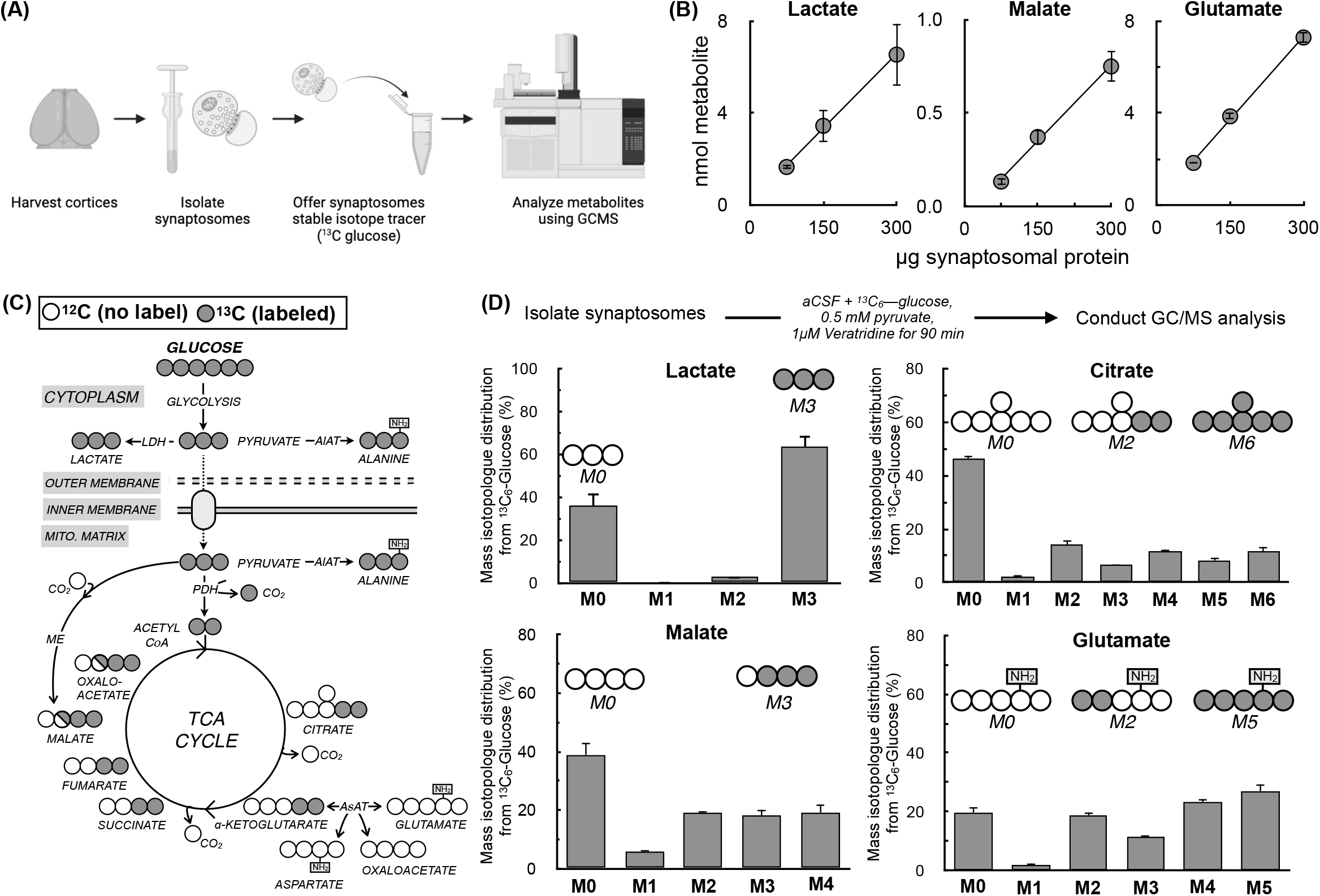
Validation of mass spectrometry and stable isotope labeling analysis of cortical synaptosomes. **(A)** Schematic outlining procedure for measuring metabolite abundance and enrichment from uniformly labeled ^13^C_6_-glucose in cortical synaptosomes. **(B)** Representative metabolites exhibit a linear relationship between signal and sample material between 75 and 300μg of synaptosomal protein. (n=3 biological replicates) **(C)** Schematic showing enrichment from ^13^C_6_-glucose into intermediary metabolism for stable isotope tracing measurements. **(D)** Mass isotopologue distributions from ^13^C_6_-glucose for representative metabolites incubated for 90 min in the presence of the 1μM veratridine. (n=3 biological replicates)

Having validated that the approach can appropriately detect metabolite abundances and isotopologue distribution patterns, we next determined whether we could reveal expected perturbations in glucose metabolism using known effector compounds with ^13^C_6_-glucose tracing. We first measured the effect of the mitochondrial uncoupler Bam15, which stimulates glucose uptake and oxidation by increasing the synaptosomal energy demand (42) (**Fig. 2A**). As expected, increased enrichment from glucose was observed in TCA cycle intermediates, the glycolytic metabolites pyruvate and lactate, and the neurotransmitters glutamate and GABA (**Fig. 2B**). We next tested the effects of UK5099, a mitochondrial pyruvate carrier (MPC) inhibitor that shifts the balance of ATP production away from oxidative phosphorylation and towards glycolysis when only glucose and pyruvate are offered as oxidizable substrates (41) (**Fig. 2C**). Indeed, UK5099 treatment reduced enrichment from labeled glucose into TCA cycle intermediates and neurotransmitters, but increased enrichment into pyruvate and lactate (**Fig. 2D**). Finally, we treated synaptosomes with the transaminase inhibitor aminooxyacetate (24) (**Fig. 2E**). As expected, enrichment patterns showed a strong reduction in enrichment of the amino acids glutamate, GABA, and alanine, while also showing a shift in the balance of energy metabolism towards glycolysis due to inhibition of the malate-aspartate shuttle (**Fig. 2F**). Altogether, the data show that stable isotope tracing in cortical synaptosomes can reliably identify perturbations in glucose metabolism.

**Figure 2.**
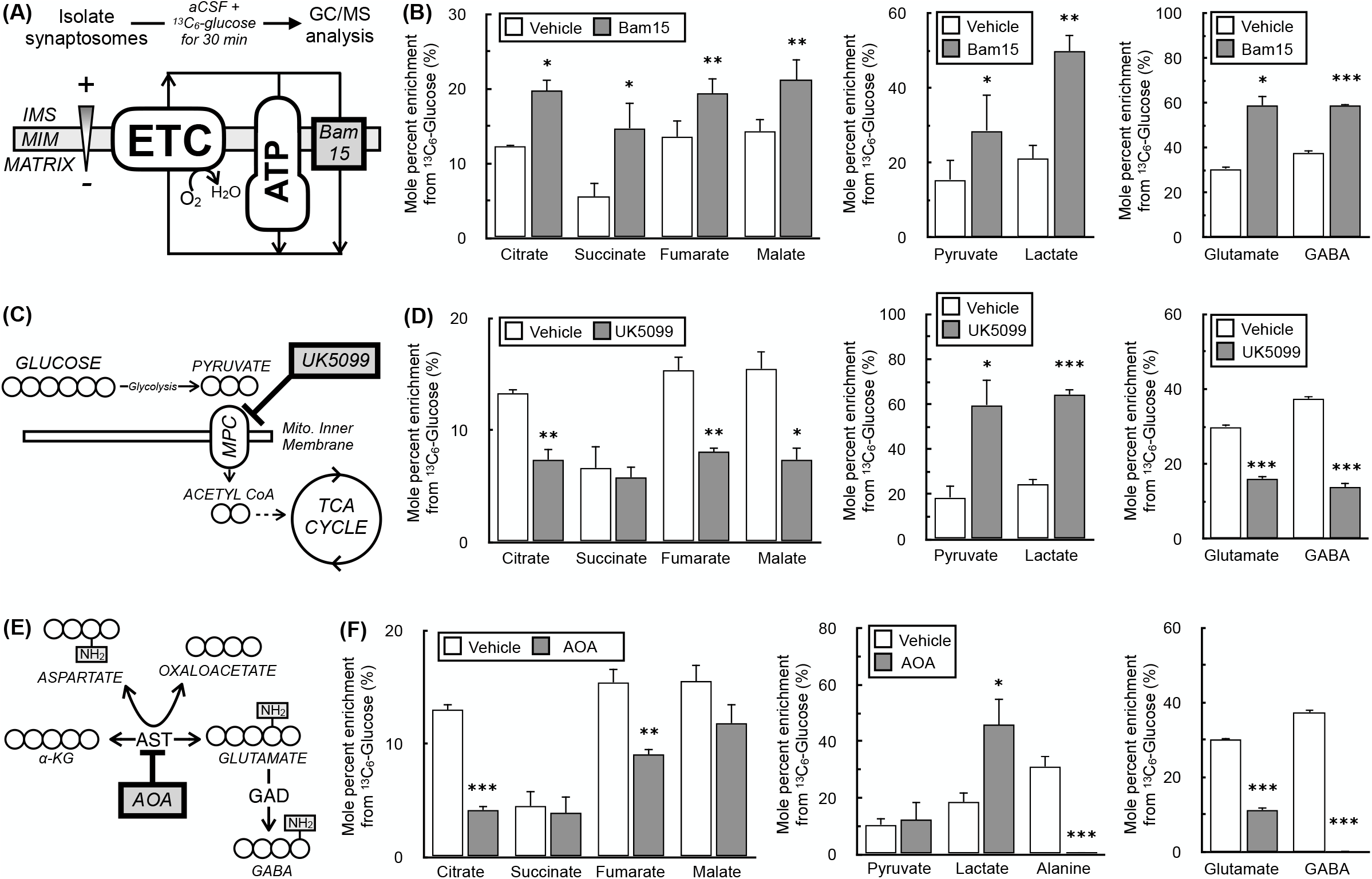
Stable isotope tracing in synaptosomes reflects expected behavior in response to metabolic perturbations. **(A)** Schematic showing the mechanism of action for the mitochondrial uncoupler Bam15. IMS, intermembrane space; MIM, mitochondrial inner membrane; ETC, electron transport chain; ATP, ATP synthase. **(B)** Mole percent enrichment (MPE) for ^13^C_6_-glucose incorporation into (*left*) TCA cycle intermediates, (*center*) glycolytic intermediates, and (*right*) glutamate and GABA in response to treatment with 50nM Bam15 for 30 min. (n=4 biological replicates) **(C)** Schematic showing the mechanism of action for the mitochondrial pyruvate carrier (MPC) inhibitor UK5099. **(D)** Mole percent enrichment (M.P.E.) for ^13^C_6_-glucose incorporation into (*left*) TCA cycle intermediates, (*center*) glycolytic intermediates, and (*right*) glutamate and GABA in response to treatment with 5μM UK5099 for 30 min. (n=4 biological replicates) **(E)** Schematic showing the mechanism of action for the transaminase inhibitor aminooxyacetate (AOA). AST, aspartate aminotransferase; GAD, glutamic acid decarboxylase. **(F)** Mole percent enrichment (MPE) for ^13^C_6_-glucose incorporation into (*left*) TCA cycle intermediates, (*center*) glycolytic intermediates and alanine, and (*right*) glutamate and GABA in response to treatment with 5μM UK5099 for 30 min. (n=4 biological replicates) The same vehicle controls were used for (B), (D), and (F) and are presented individually against a given effector for clarity. All data are presented as mean ± standard error of the mean (S.E.M.) of individual biological replicates. Statistical analysis was conducted with a pairwise Student’s t test for each individual metabolite for a given treatment against vehicle control. *, p < 0.05; **, p < 0.01; ***, p < 0.001.

### Method Application: 5X-FAD mice treated with Dapagliflozin

To apply the method to better understand the links between neuronal metabolic dysfunction and cognitive impairment, we measured synaptosomal glucose metabolism and bioenergetics in the 5X-FAD mouse model of AD. Given the links between AD and insulin resistance, we further tested the effect of the SGLT-2 inhibitor Dapagliflozin on both groups. SGLT-2 inhibitors are widely prescribed antidiabetic drugs that block the renal reabsorption of glucose resulting in urinary glucose loss that promotes euglycemia in people with diabetes (43), and have been recently shown to have remarkable cardioprotective effects in patients independently from lowering blood glucose (32, 44). Dapagliflozin was administered to animals at 10mg/kg via the drinking water for 8 weeks prior to behavioral testing and euthanasia. Given the sex differences in the 5X-FAD model – females can exhibit amyloid burden and cognitive deficits earlier than males (34) – female animals were treated beginning at 5 months of age (sacrificed at 7 months) whereas males were treated beginning at 6 months of age (sacrificed at 8 months).

Urinary glucose measurements showed robust target engagement, as Dapagliflozin increased glycosuria by over three orders of magnitude regardless of transgene status or sex (**Figs. 3A&B**). Unexpectedly, the 5X-FAD animals also had lower unfasted plasma glucose levels than control animals, and treatment with Dapagliflozin trended towards restoring plasma glucose to control levels (**Fig. 3C**). This effect was largely driven by the male cohort (**Fig. 3D**). Finally, we measured SGLT2 protein content in the cerebral cortex and kidney of mice to confirm any effects observed from SGLT2 inhibition were primarily on peripheral metabolism rather than a direct neural mechanism of action. As expected, SGLT2 was undetectable in the brain but highly expressed in the kidney (**Fig. 3E**). Altogether, the data demonstrate successful renal SGLT2 inhibition during the 8 weeks of Dapagliflozin treatment.

**Figure 3.**
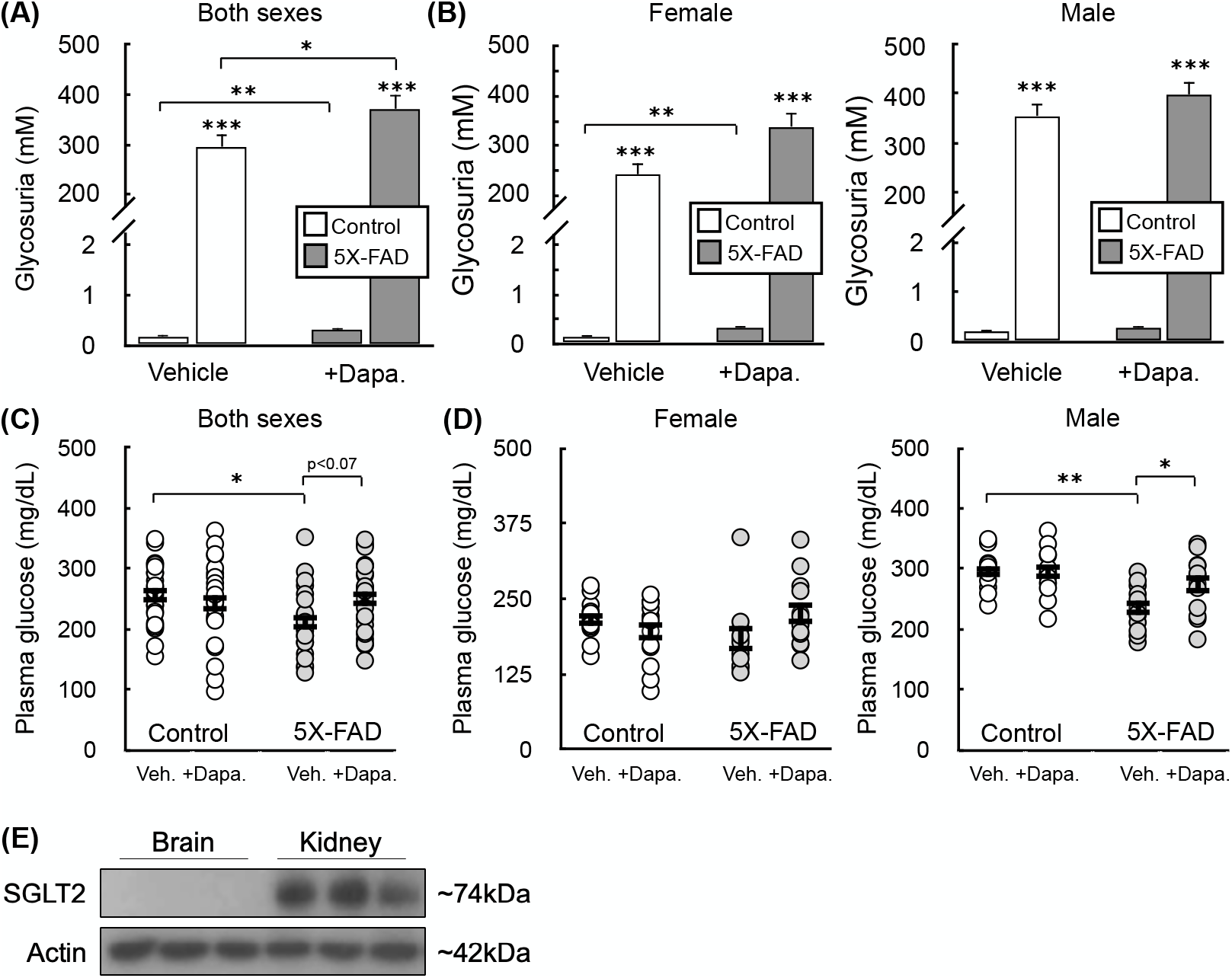
Target engagement upon two-month Dapagliflozin treatment. **(A)** Urinary glucose measured in WT and 5X-FAD animals after two months of treatment with Dapagliflozin (10 mg/kg) administered via the drinking water. (n=22 animals). **(B)** Urinary glucose as in (A) separated by sex (n ≥ 10 animals). **(C)** Plasma glucose measured in WT and 5X-FAD animals after two months of treatment with Dapagliflozin (10 mg/kg) administered via the drinking water. Animals were not fasted prior to measurements (n=22 animals). **(D)** Plasma glucose as in (C) separated by sex (n≥10 animals). **(E)** Western analysis of relative SGLT2 levels in the brain and kidney (n=3 animals). All data are presented as mean ± standard error of the mean (S.E.M.) of individual biological replicates. Statistical analysis was conducted with analysis of variance (ANOVA) with Dunnett’s *post hoc* test. *, p < 0.05; **, p < 0.01; ***, p < 0.001.

After confirming target engagement, we measured short-term, spatial working memory with the Y-maze test in control and 5X-FAD animals. The test consists of repeatedly placing animals in a three-armed structure in the shape of a ‘Y,’ and is based on the predisposition of rodents to explore a new arm of the maze rather than return to the one it previously visited (35). We chose the 5-month (female) and 6-month (male) time point to begin Dapagliflozin treatment in order to intervene relatively early and observe any putative delay in the onset of pathology. Indeed, prior to sacrifice at 7 (female) or 8 months (male), 5X-FAD animals showed a reduced propensity to explore new arms of the maze (**Fig. 4A**, *left*) as well as a trend towards lower activity (**Fig. 4A**, *right*). Separating the animals by sex revealed the memory impairments were largely observed in the female cohort (**Fig. 4B**). Moreover, examining the effect of SGLT-2 inhibition showed a trend towards improved Y-maze performance in the female 5X-FAD animals treated with Dapagliflozin. We then conducted immunohistochemistry on WT and 5X-FAD female mice to visually confirm that amyloidosis had developed at the time of sacrifice. We subsequently followed up with an ELISA-based assay for soluble β-amyloid 1-42 in both the hippocampus (**Fig. 4D**) and cortex (**Fig. 4E**) of 5X-FAD animals treated with Dapagliflozin. Dapagliflozin did not have any effect on amyloid burden in the hippocampus, and sex effects could not be measured due to an insufficient number of samples. In the cortex, however, male animals developed a more pronounced amyloid burden which was significantly reduced by SGLT2 inhibition, whereas drug treatment had no effect on the lower amyloid levels detected in females. The data show that Dapagliflozin had a mild effect on preserving transgene-induced spatial memory deficits (in females) and reducing amyloid burden (in males).

**Figure 4.**
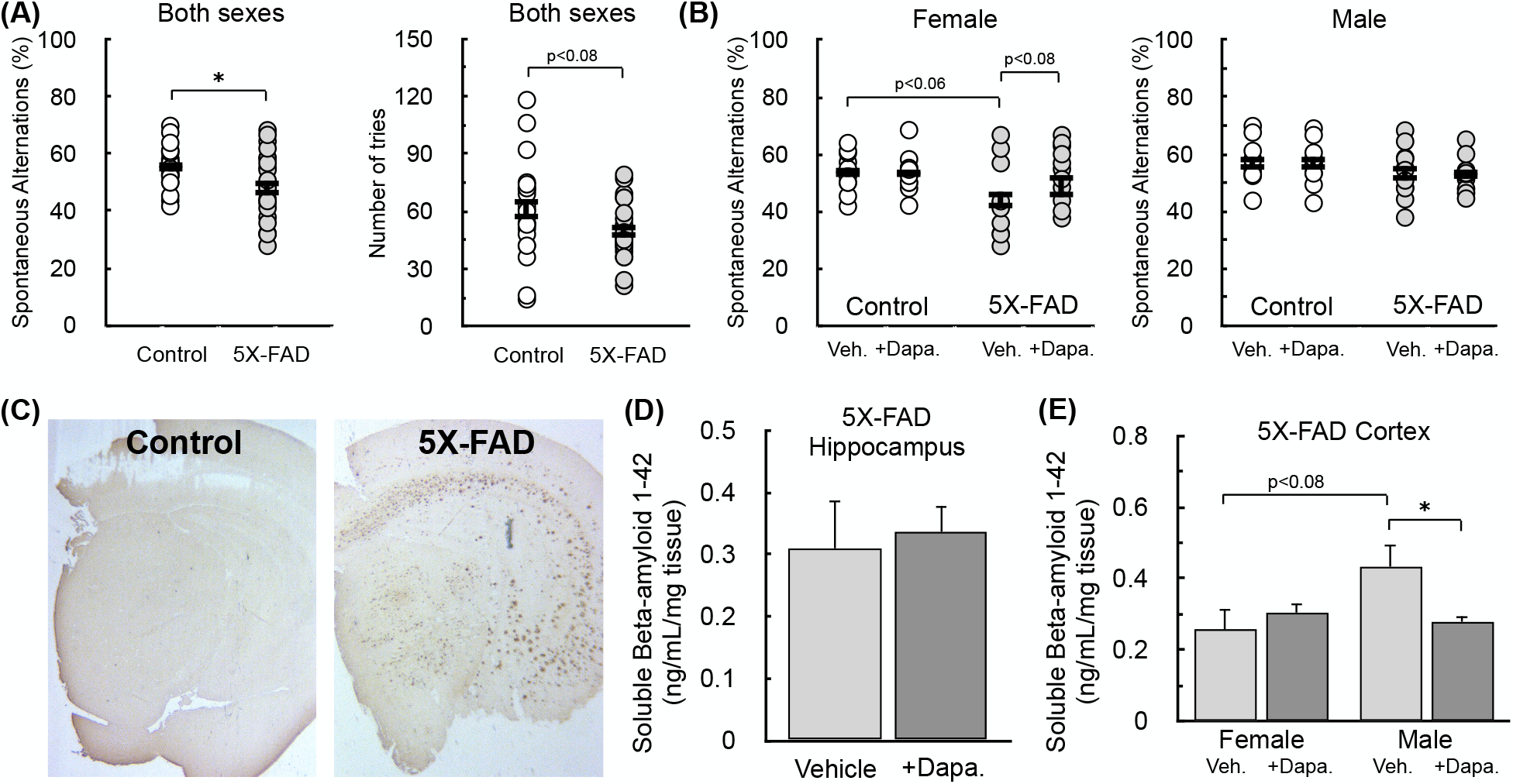
Behavioral analysis and amyloidosis following two-month Dapagliflozin treatment. **(A)** (*Left*) Spontaneous alternations and (*right*) number of tries in the Y-maze spatial memory test. (n=22 animals) from control and 5X-FAD animals treated with either 10 mg/kg Dapagliflozin or vehicle controls. **(B)** Spontaneous alternations as in (A) separated by sex (n≥10 animals). **(C)** Representative immunohistochemistry image at the time of sacrifice from both a control and 5X-FAD female animal stained with a polyclonal antibody against the first 13 amino acids of β-amyloid. **(D-E)** ELISA-based measurements against β-amyloid residues 1-42 in the TBS-soluble fraction from the (D) cortex and (E) hippocampus of 5X-FAD mice treated with either 10 mg/kg Dapagliflozin or vehicle controls. Results from the cortex were separated by sex, but insufficient samples were available to do so for the hippocampus. (n 6 animals). All data are presented as mean ± standard error of the mean (S.E.M.) of individual biological replicates. Statistical analysis was conducted with either a Student’s t test for pairwise comparison or analysis of variance (ANOVA) with Dunnett’s *post hoc* test as appropriate. *, p < 0.05; **, p < 0.01; ***, p < 0.001.

We next sought to examine bioenergetics from the cortical synaptosomes harvested from these animals. We first conducted respirometry on permeabilized synaptosomes. Permeabilizing the plasma membrane of synaptosomes with recombinant perfringolysin O (rPFO) enables studies of pathway-specific respirometry normally done with isolated mitochondria to be conducted on cells and synaptosomes (45), from which mitochondrial isolation is usually prohibited by small sample size. Synaptosomes were offered various substrates to test specific mitochondrial oxidative pathways (**Fig. 5A**): pyruvate with malate (**Fig. 5B**), glutamate with malate (**Fig. 5C**), β-hydroxybutyrate with malate (**Fig. 5D**), and succinate with rotenone (**Fig. 5E**). No differences were observed with either transgene status or Dapagliflozin treatment across either sex (aggregate data for both sexes presented here). Nonetheless, the absence of an effect in this reductionist approach – which measures maximal pathway activities conducted under non-physiological conditions – does not necessarily indicate a lack of metabolic or even mitochondrial defects (14).

**Figure 5.**
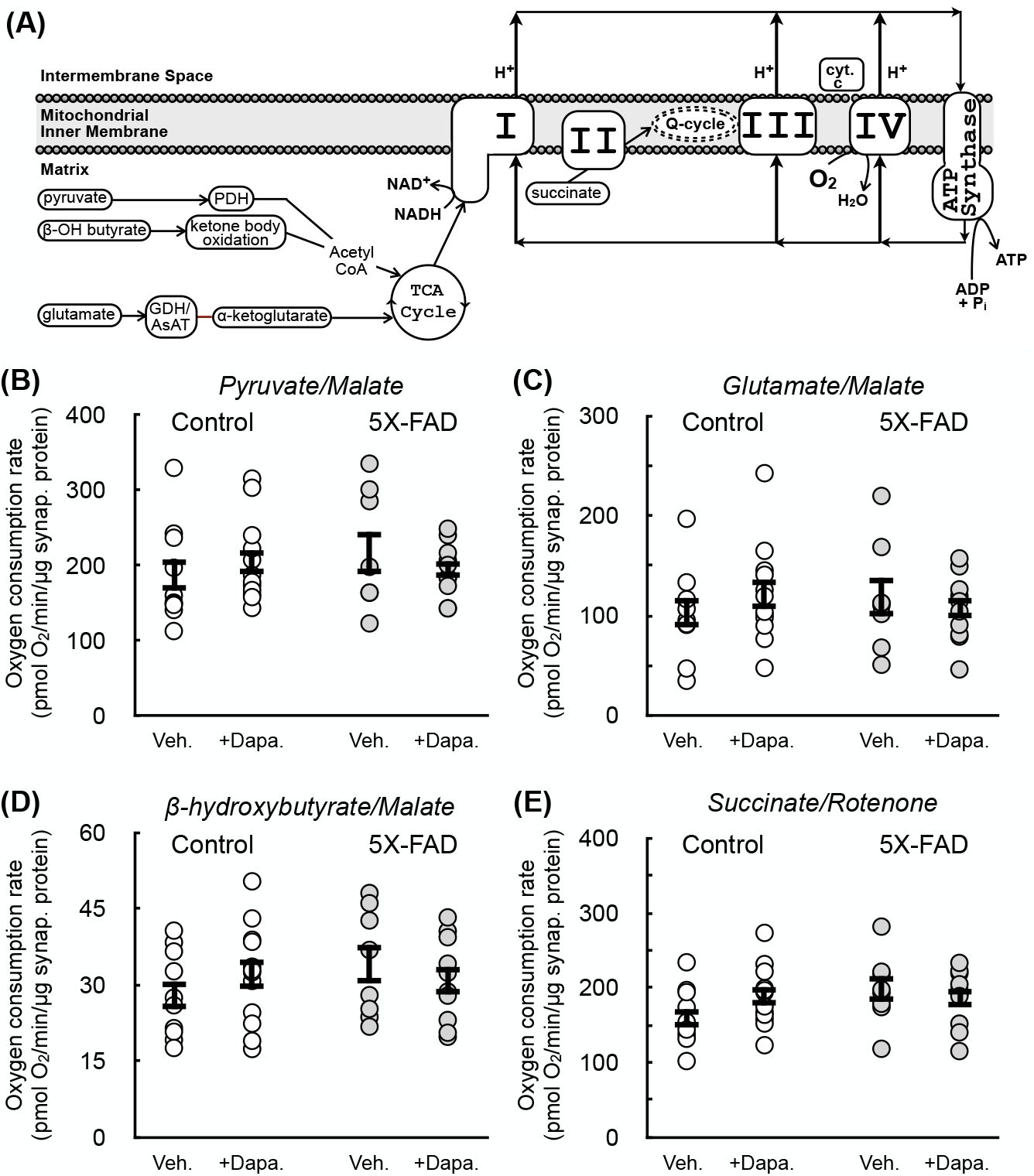
No change observed with respirometry in permeabilized synaptosomes between control and 5X-FAD animals. **(A)** Scheme detailing how various respiratory substrates feed into oxidative phosphorylation. PDH, pyruvate dehydrogenase; GDH, glutamate dehydrogenase; AsAT, aspartate aminotransferase, cyt. c, cytochrome c. **(B-E)** ADP-stimulated (“State 3”) respiration in rPFO-permeabilized cortical synaptosomes from WT and 5X-FAD animals treated with either 10 mg/kg Dapagliflozin or vehicle controls. Permeabilized synaptosomes were offered as respiratory substrates: (B) pyruvate with malate, (C) Reglutamate with malate, (D) β-hydroxybutyrate with malate, or (E) succinate with rotenone (n≥10 biological replicates of mixed sex). All data are presented as mean ± standard error of the mean (S.E.M.) of individual biological replicates. Statistical analysis was conducted with analysis of variance (ANOVA) with Dunnett’s *post hoc* test.

We therefore harvested cortical synaptosomes from the female cohort of animals for stable isotope tracing with uniformly labeled ^13^C_6_-glucose. These animals showed trends towards reduced short-term spatial memory with 5X-FAD status and improved performance with Dapagliflozin treatment. Indeed, mass spectrometry results showed changes with both 5X-FAD status and SGLT-2 inhibition that were associated with results from the Y-maze test. Levels of TCA cycle intermediates, glutamate, and aspartate all showed a trend towards reduced levels in synaptosomes harvested from 5X-FAD, and a restoration of levels in animals treated with Dapagliflozin (**Figs. 6A&B**; data for all metabolites analyzed and separated by sex is given in **Supplemental Table 1**). Metabolite levels harvested from the retrosplenial cortex, however, were unchanged regardless of transgene or treatment status, underlying the importance of regional and cell-type heterogeneity of brain energy metabolism (**Supplemental Table 2**). Stable isotope tracing also revealed specific metabolic defects in glucose metabolism with 5X-FAD status. Incorporation of glucose-derived carbon into citrate was significantly lower in synaptosomes harvested from 5X-FAD animals, and this loss was almost entirely prevented with Dapagliflozin treatment (**Figs. 6C&D**). However, no changes in enrichment were observed in pyruvate, lactate, or malate (**Fig. 6E**), suggesting a specific change in pyruvate uptake and/or oxidation. We therefore examined the phosphorylation status of pyruvate dehydrogenase (PDH) in the TBS-soluble fraction of bulk cortex from female animals. Unexpectedly, however, we observed a trend towards reduced PDH phosphorylation in 5X-FAD animals (**Fig. 6F**), which is usually associated with increased activity as the PDH kinases are inhibitory (46). Altogether, though, the results show that metabolomics and ^13^C stable isotope tracing in isolated synaptosomes can pinpoint changes in glucose metabolism difficult or impossible to obtain with other approaches.

**Figure 6.**
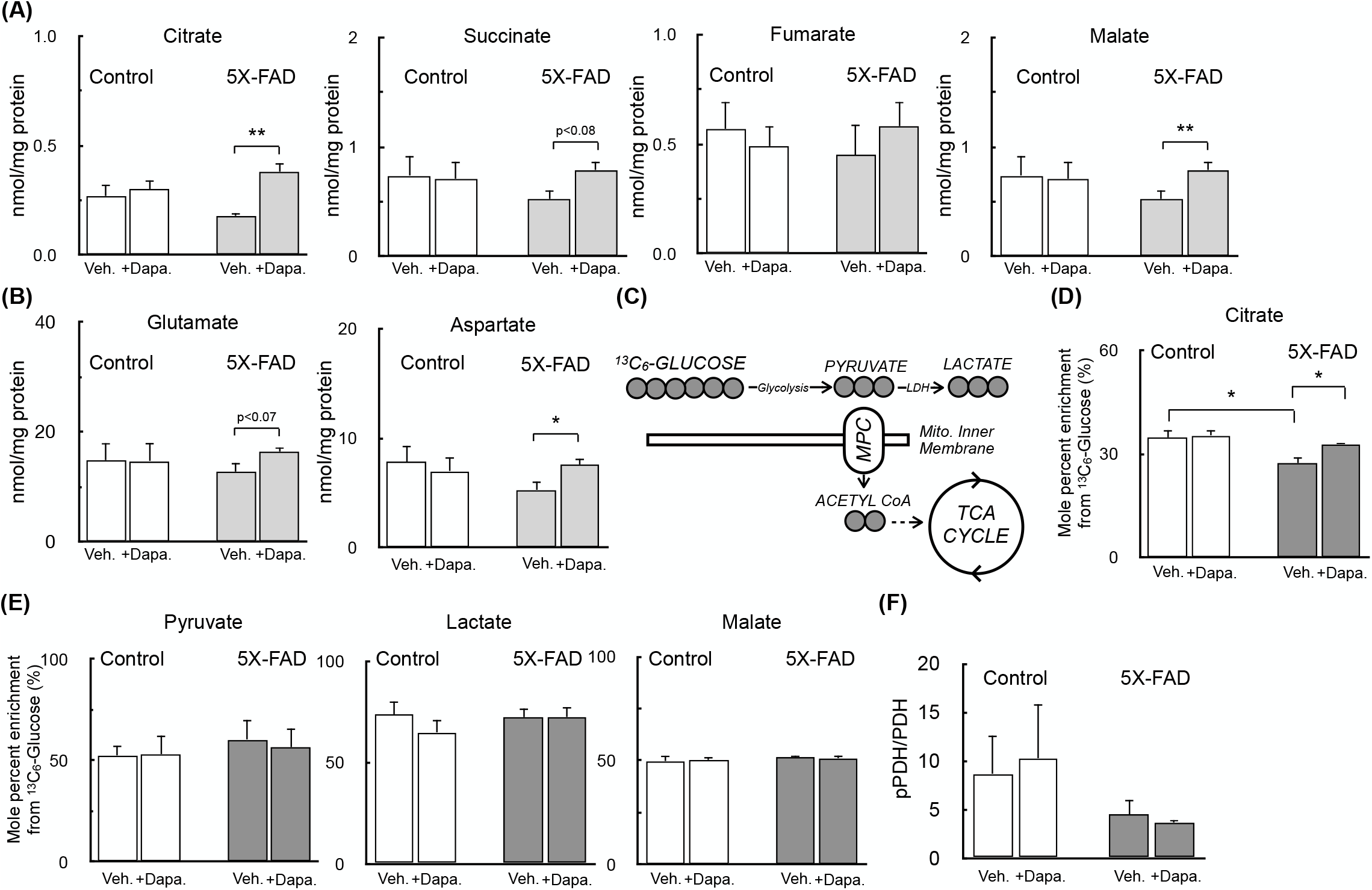
Preserved synaptosomal metabolite abundance and pyruvate oxidation in 5X-FAD female mice with Dapagliflozin treatment. **(A)** Metabolite abundance of for TCA cycle intermediates in cortical synaptosomes assayed as in 1D from female control and 5X-FAD animals treated with either 10 mg/kg Dapagliflozin or vehicle controls. (n 4≥biological replicates) **(B)** Abundance of glutamate and aspartate for synaptosomes as in (A). (n≥4 biological replicates) **(C)** Schematic showing enrichment from ^13^C_6_-glucose into intermediary metabolism for stable isotope tracing measurements. **(D&E)** Mole percent enrichment (MPE) for ^13^C_6_-glucose incorporation into (D) citrate and (E) pyruvate, lactate, and malate for synaptosomes treated as in (A). (n≥4 biological replicates) **(F)** Quantitative western analysis for to examine the phosphorylation state of pyruvate dehydrogenase. (n≥4 biological replicates) in the TBS-soluble fraction of the bulk cortex from female animals. All data are presented as mean ± standard error of the mean (S.E.M.) of individual biological replicates. Statistical analysis was conducted with analysis of variance (ANOVA) with Dunnett’s *post hoc* test.

## DISCUSSION

Stable isotope tracing is a powerful technique that can reveal metabolic fluxes through individual enzymes and pathways under native, basal conditions. As such, it can reveal metabolic dysfunction not possible with destructive endpoint measurements (e.g. gene or protein expression, enzymology) or reductionist bioenergetic approaches with isolated mitochondria. Here we demonstrate that this technique is suitable to study metabolism in isolated synaptic nerve terminals, with validation experiments able to detect specific metabolic perturbations such as an increased energy demand or targeted enzyme inhibition. Moreover, the labeling patterns themselves can reveal important information about neuronal metabolism given the relative purity of the preparation (47). Although it is widely thought that astrocytes are almost exclusively responsible for glutamate synthesis in the brain (48, 49), the relatively high proportion of M+3 and M+5 labeling from ^13^C_6_-glucose into glutamate and TCA cycle intermediates suggest a high degree of glucose anaplerosis in neurons and, specifically, synaptic nerve terminals (41, 50).

Our GC/MS analysis of synaptosomes isolated from 5X-FAD animals treated with Dapagliflozin further highlights the strength of this technique. Specifically, the reduced glucose enrichment into citrate and steady-state TCA cycle intermediates levels in synaptosomes from female 5X-FAD animals – and their restoration by Dapagliflozin – could not have been identified by other common techniques. For example, measurement of maximal pyruvate oxidation in permeabilized synaptosomes from the same cohort of animals showed no change with either 5X-FAD status or Dapagliflozin treatment, perhaps suggesting a level of regulation not easily identified with reductionist approaches (e.g. posttranslational modification of pyruvate transport or oxidation). Moreover, western analysis of PDH phosphorylation status from whole cortex showed no change with Dapagliflozin treatment, perhaps indicating a neuronal-specific effect lost in bulk tissue or a change in mitochondrial pyruvate uptake rather than oxidation. Nonetheless, mitochondrial pyruvate oxidation has been identified by several reports as an important regulatory node of metabolic dysfunction in Alzheimer’s Disease (51–54), a finding reinforced by our stable isotope tracing data.

Our data showed these changes in synaptosomal metabolism in female mice were associated with improved short-term spatial working memory. Findings with the Y-maze test have been mixed in the 5X-FAD model (55–57) and statistical analysis for this test may be uniquely sensitive to effects of small sample size relative to other tests for cognition and locomotive behavior (58). Here we present a trend towards reduced performance associated with 5X-FAD status in females (at 8 months) with no change in males (at 9 months). Indeed, SGLT-2 inhibition showed a female-specific effect as well, though there was no observable deficit to rescue in male animals. Our study sought to intervene relatively early in our animal cohort to test whether Dapagliflozin could delay the development of pathology, as opposed to treating older animals to examine the effect of SGLT-2 inhibition on established disease. This design, along with a small sample size, are likely factors in explaining the small, subtle nature of the observed changes.

Unexpectedly, the trends in Y-maze performance were not associated with amyloid burden in the TBS-soluble fraction collected from either the hippocampus or cortex. In fact, cortical samples from the male cohort – harvested one month later than the females – showed a significant reduction in β-amyloid after Dapagliflozin treatment. No significant effects were observed in the female cohort, probably explained by the lower amyloid burden in females compared to males. Moreover, it is important to note that the amyloid measurements were conducted in bulk tissue and may not reflect localized or cell type-specific changes. Although we did not find that amyloid deposition correlated with cognitive impairments, the results reinforce that SGLT-2 inhibition may be an attractive mechanism to improve cognition (31, 59, 60) as well as proteostasis under conditions of a high amyloid burden (33, 61).

## EXPERIMENTAL PROCEDURES

### Animals

All animal experiments were performed in accordance with the National Institutes of Health regulations, approved by the UCLA Animal Research Committee, and performed under oversight of the Division of Laboratory Animal Medicine. 5X-FAD animals (006554; MMRRC Strain #034840-JAX) and controls (100012 B6SJLF1/J) were purchased from Jackson Laboratories before 5 months of age. Animals were housed in groups of up to four in individually ventilated cages under standard conditions (22°C, 12-hour light-dark cycle) receiving food and water *ad libitum*.

Beginning at age 5 months (females) or 6 months (males) animals were randomly split into treatment groups and offered Dapagliflozin at 10 mg/kg via the drinking water with bottles wrapped in aluminum foil to protect against light-induced damage. Dapagliflozin was administered for 8 weeks, after which time animals were sacrificed. Different sexes were sacrificed at different ages in order to account for the heavier amyloid burden in females in the 5X-FAD model (34). Glucose in the urine and plasma was measured using commercial test strips (NovaMax). Animals were not fasted prior to measurements.

Short-term spatial working memory was assessed using the Y-maze test (35). Animals were allowed to freely explore in the maze for 8 min. The box was illuminated with a 60W overhead light bulb and animals were filmed with an overhead camera. AnyMaze video tracking software was used to analyze movements.

### Isolation of cortical synaptic nerve terminals (‘synaptosomes’)

Isolated synaptosomes from mouse cortices were isolated by a Percoll gradient separation and centrifugation broadly in line with standard procedures (21, 36). All steps were conducted on ice or at 4°C. Isolated cortices were diced and homogenized using a glass-on-glass 7mL Dounce homogenizer (Wheaton) using 10 strokes of a loose pestle followed by 5-10 strokes of a tight pestle in medium containing 320mM sucrose, 1mM EDTA, and 0.25mM dithiothreitol (DTT). The homogenate was centrifuged in 1.5 mL microfuge tubes at 1,300x*g* for 10 min, and the supernatant laid at the top of a Percoll gradient in 16 mL ultracentrifuge tubes. The gradient consisted of 3 layers – 23%, 10%, and 3% (all v/v in the sucrose, EDTA, DTT solution as described earlier). Tubes were centrifuged for 8 min at 32,500x*g* to enrich the synaptosomes between the 23% and 10% Percoll solutions. This layer was removed using a serological pipette and resuspended in 6-10 mL of Synaptosomal Resuspension Medium with the following composition: 140mM NaCl, 20 mM HEPES (pH 7.4), 5mM KCl, 5 mM NaHCO_3_, 1mM MgCl_2_, 1.2 mM Na_2_HPO_4_, and 10 mM glucose. This resuspension was then spun at 20,000x*g* for 18 min to isolate the synaptosomes and remove residual Percoll. The supernatant was aspirated and the loose pellet was resuspended to a final volume of ∼200μL in Synaptosomal Resuspension Medium. For studies involving 5X-FAD animals and controls, the brains of anaesthetized animals were perfused with 30 mL of perfusion buffer via the left ventricle according to standard procedures. The perfusion buffer consisted of the following: 137 mM NaCl, 10 mM HEPES, 4.6 mM KCl, 1.1 mM K_2_HPO_4_, 1.1 mM EDTA, 0.6 mM MgSO_4_, protease inhibitors (1μg/mL leupeptin, 1μg/mL aprotinin, 1μg/mL pepstatin, and 50μg/mL PMSF), and phosphatase inhibitors (1mM sodium vanadate, 1mM sodium pyrophosphate, and 20mM sodium fluoride). Synaptosomal protein content was measured by the bicinconinic acid (BCA) assay.

### GC/MS and stable isotope tracing

150μg of synaptosomes were resuspended in microfuge tubes containing 1mL of artificial cerebrospinal fluid (aCSF: 120 mM NaCl, 3.5 mM KCl, 1.3 mM CaCl_2_, 0.04 mM KH_2_PO_4_, 1 mM MgCl_2_, 5 mM HEPES, and 3μg/L phenol red) supplemented with 10 mM uniformly labeled U-^13^C_6_ glucose, 0.5 mM pyruvate, and 1μM veratridine at 37°C for either 30 or 90 min as indicated for a given experiment. Veratridine was omitted from the experimental medium when assessing the response to mitochondrial effector compounds in Fig. 2. Samples were centrifuged at 4°C for 5 min at 3,000x*g*. Samples were then washed with ice cold 0.9% (w/v) saline and centrifuged again at 4°C for 5 min at 3,000x*g*. Polar metabolites were extracted with a Folch-like method with 500μL methanol, 200μL water containing 1μg norvaline as an internal standard, and 500 μL chloroform. Samples were vortexed and centrifuged at 4°C for 5 min at 10,000x*g* and the polar, top layer was collected and dried using a refrigerated centrivap. Polar metabolites were derivatized and analyzed via GC/MS analysis according to well established methods (23, 37). For targeted detection of metabolites in the retrosplenial cortex, the tissue was snap-frozen in liquid nitrogen, weighed, and disrupted with a bead-based tissue lyser (Qiagen) in the Folch-like extraction solution. Metabolite levels were quantified against a standard curve of quantities ranging from 70 pg to 50 ng of each metabolite.

### Respirometry

Oxygen consumption was measured in permeabilized synaptosomes using a Seahorse XF96 Analyzer in line with previously established protocols (12). Synaptosomes were seeded onto poly-D-lysine-coated Seahorse XF96 plates at either 2 μg/well (complex I-linked substrates) or 1 μg/well (succinate/rotenone). Synaptosomes were assayed in MAS medium [220mM mannitol, 70mM sucrose, 10mM KH_2_PO_4_, 5mM MgCl_2_, 1mM EGTA, and 2mM HEPES (pH 7.2)] supplemented with 4mM ADP, 0.2% (w/v) de-fatted BSA, 3nM recombinant perfringolysin O (rPFO), and respiratory substrates. Permeabilized synaptosomes were offered the following respiratory substrates: 5mM pyruvate with 1mM malate, 5mM glutamate with 5 mM malate, 5 mM β-hydroxybutyrate with 1 mM malate, or 2 mM succinate with 2μM rotenone. For complex I-linked substrates, 1 mM NAD^+^ was added to the assay medium (38). ADP-sensitive respiration (‘State 3’) was measured and corrected for background signal with 200nM rotenone and 1μM antimycin A (14).

### β-amyloid measurements

For immunohistochemistry, formalin-fixed, paraffin-embedded mouse hemi-brains were sectioned into 10 μm slices and mounted onto slides. Slides were de-paraffinized and quenched using 0.3% (v/v) hydrogen peroxide in methanol for 35 minutes. After pre-treatment with formic acid for 10 min at room temperature, sections were treated with a blocking solution that consisted of 5% (v/v) normal goat serum in 3% (v/v) bovine serum albumin (BSA) for 45 minutes at 37°C. Sections were then incubated with the primary antibody (anti-’DAE’, 1:200 dilution) for 1 hour at 37°C, then overnight at 4°C. The ‘DAE’ antibody is an N-terminal peptide developed in rabbits against residue 1-13 of β-amyloid (39). The following day, the sections were incubated with the secondary antibody (anti-rabbit, 1:1000 dilution) in 1.5% (v/v) normal goat serum with 3% (v/v) BSA. Endogenous biotin and avidin was blocked using a Avidin/Biotin Blocking Kit (Vector Laboratories; Burlingame, CA) for 75 min at 37°C. The slides were then visualized using 3,3’-diaminobenzidine (DAB). Images of brain sections were taken at 2x magnification with a digital MC170 5 MPixel Leica camera on an Olympus Vannox-T (AHBT) microscope. The Bregma coordinate of sections was -2.06mm.

For ELISA-based measurements, cortices and hippocampi were snap-frozen in liquid nitrogen and later weighed and extracted in three phases. First, soluble proteins were extracted into supernatant by adding 10X Tris-buffered saline (TBS) for each the volume of the brain weight containing complete protease inhibitor (PI) and phosphatase inhibitor (PPI) cocktail (Roche) before sonication for three 10 sec pluses on ice. The samples were then centrifuged at 132,000x*g* for 20 min at 4ºC and the supernatant collected into a separate tube. RIPA lysis buffer containing PI and PPI was then added to the leftover insoluble pellet for the second extraction, followed by sonication, centrifugation at 132,000x*g* for 20 min at 4ºC, and supernatant collection. Finally, a Tris-HCl solution containing 6.25M guanidine was added to the leftover pellet to extract ß-amyloid. The samples were then sonicated and centrifuged. The supernatant from the soluble fraction was collected for use with the V-PLEX Human Aß42 Kit (MSD, Rockville, MD) to quantify β-amyloid 1-42.

### Western analysis

Mouse cortices from perfused brains as described above were sonicated in Tris-buffered saline (TBS) containing protease and phosphatase inhibitor cocktails previously developed (40). The samples were then centrifuged at 100,000x*g* for 20 min and the supernatant retained to obtain a soluble fraction. Protein content was determined using a BCA protein assay kit, and 30μg of protein resuspended in Laemmli buffer was loaded into each well of a 7-20% Tris-glycine gradient gel. Proteins were then transferred to PVDF membranes, blocked with 5% nonfat milk in TBS with 0.1% Tween-20 for 1 hr. Samples were assessed for the fraction of phosphorylated PDH (Abcam ab#31866S; 1:1,000) relative to total protein (ab#3205; 1:1,000). An HRP-conjugated secondary antibody (1:5000; Thermo Fisher) was used for densitometry quantification by Image J, and β-Actin was used as a loading control (sc-47778, 1:2,000 Santa Cruz Biotechnology). For detection of SGLT-2 in the brain and kidney, tissue samples were disrupted in RIPA buffer [50 mM Tris, 150 nM NaCl, 0.1%(w/v) SDS, 0.5%(w/v) sodium deoxycholate, 1%(v/v) NP-40, and 5 mM EDTA], solubilized in Laemmli buffer and heated to 95°C for 5 min, and loaded at 50μg/well for western as before [Abcam ab85626 (1:500)].

### Statistical Analysis

All statistical parameters, including the number of biological replicates (n), can be found in the figure legends. Statistical analyses were performed using Graph Pad Prism 10 software. Data are presented as the mean ± standard error of the mean (SEM). Individual pairwise comparisons were performed using two-tailed Student’s t-test. For experiments involving two or more factors, data were analyzed by one-way, repeated measures ANOVA followed by Tukey’s *post hoc* multiple comparisons tests. Data were assumed to follow a normal distribution (no tests were performed). Values denoted as follows were considered significant: *p < 0.05; **p <0.002; ***p<0.001.

## DATA AVAILABILITY

All data supporting the findings in this study are available from the corresponding author upon request.

## ACKNOWLEDGEMENTS

This work was supported by AstraZeneca (NCR-19-145) to ASD, GMC, and OSS. ASD is also supported by National Institutes of Health (NIH) Grant R35GM138003 as well as the W.M. Keck Foundation (995337) and the Agilent Early Career Professor Award (4818). SAF is supported by R01AG066212. MV was supported by the UCLA Chemistry-Biology Interface Training Grant (T32GM136614).

