## Supplemental Table 1 for "^13^C tracing in synaptosomes reveals that SGLT2 inhibition with dapagliflozin prevents metabolic deficits in the 5X-FAD model of Alzheimer’s Disease"

**Supplemental Table 1**: Metabolite abundances in cortical synaptosomes acutely offered glucose and pyruvate

| | BOTH SEXES (nmol/mg protein) | | | |
|---|---|---|---|---|
| | WT | | 5X-FAD Control | |
| | Control | DAPA | Control | DAPA |
| **ALANINE** | **0.83** | **0.83** | **0.99** | **0.94** |
| S.E.M. | 0.09 | 0.06 | 0.10 | 0.07 |
| **ASPARTATE** | **12.30** | **13.69** | **9.84** | **14.86** |
| S.E.M. | 2.43 | 2.52 | 2.20 | 3.14 |
| **CITRATE** | **0.43** | **0.68** | **0.38** | **0.66** |
| S.E.M. | 0.10 | 0.13 | 0.09 | 0.12 |
| **FUMARATE** | **0.87** | **0.98** | **0.80** | **0.84** |
| S.E.M. | 0.17 | 0.16 | 0.17 | 0.13 |
| **GLUTAMATE** | **16.66** | **19.57** | **17.81** | **19.50** |
| S.E.M. | 2.14 | 2.77 | 2.16 | 1.75 |
| **LACTATE** | **15.19** | **14.22** | **14.04** | **13.94** |
| S.E.M. | 2.21 | 2.01 | 2.44 | 2.75 |
| **MALATE** | **3.83** | **4.68** | **3.12** | **4.24** |
| S.E.M. | 0.85 | 0.77 | 0.85 | 0.74 |
| **PYRUVATE** | **2.95** | **3.31** | **3.25** | **2.94** |
| S.E.M. | 0.41 | 0.66 | 0.51 | 0.42 |
| **SERINE** | **0.79** | **0.71** | **0.63** | **0.91** |
| S.E.M. | 0.17 | 0.11 | 0.08 | 0.12 |
| **SUCCINATE** | **0.99** | **1.00** | **0.89** | **1.22** |
| S.E.M. | 0.15 | 0.12 | 0.15 | 0.19 |

| | FEMALE (nmol/mg protein) | | | |
|---|---|---|---|---|
| | WT | | 5X-FAD Control | |
| | Control | DAPA | Control | DAPA |
| **ALANINE** | **0.75** | **0.74** | **0.94** | **0.93** |
| S.E.M. | 0.10 | 0.11 | 0.17 | 0.08 |
| **ASPARTATE** | **7.82** | **7.00** | **5.28** | **7.58** |
| S.E.M. | 1.52 | 1.26 | 0.77 | 0.59 |
| **CITRATE** | **0.27** | **0.30** | **0.18** | **0.38 *** |
| S.E.M. | 0.05 | 0.04 | 0.01 | 0.04 |
| **FUMARATE** | **0.57** | **0.49** | **0.45** | **0.58** |
| S.E.M. | 0.12 | 0.09 | 0.14 | 0.11 |
| **GLUTAMATE** | **14.76** | **14.71** | **12.64** | **16.35** |
| S.E.M. | 3.10 | 3.26 | 1.78 | 0.72 |
| **LACTATE** | **15.86** | **14.99** | **17.91** | **20.72** |
| S.E.M. | 1.94 | 2.70 | 3.29 | 2.68 |
| **MALATE** | **2.42** | **2.12** | **1.25** | **2.45 *** |
| S.E.M. | 0.59 | 0.32 | 0.26 | 0.26 |
| **PYRUVATE** | **2.54** | **3.37** | **3.57** | **2.86** |
| S.E.M. | 0.51 | 1.39 | 0.98 | 0.61 |
| **SERINE** | **0.47** | **0.44** | **0.52** | **0.83** |
| S.E.M. | 0.09 | 0.14 | 0.12 | 0.21 |
| **SUCCINATE** | **0.74** | **0.71** | **0.52** | **0.79** |
| S.E.M. | 0.18 | 0.16 | 0.08 | 0.08 |

| | MALE (nmol/mg protein) | | | |
|---|---|---|---|---|
| | WT | | 5X-FAD Control | |
| | Control | DAPA | Control | DAPA |
| **ALANINE** | **0.92** | **0.91** | **1.04** | **0.95** |
| S.E.M. | 0.15 | 0.07 | 0.12 | 0.12 |
| **ASPARTATE** | **16.79** | **19.26** | **14.40** | **22.15** |
| S.E.M. | 3.78 | 2.93 | 2.86 | 4.19 |
| **CITRATE** | **0.59** | **1.00** | **0.57** | **0.94** |
| S.E.M. | 0.17 | 0.14 | 0.10 | 0.14 |
| **FUMARATE** | **1.18** | **1.38** | **1.15** | **1.09** |
| S.E.M. | 0.28 | 0.13 | 0.18 | 0.18 |
| **GLUTAMATE** | **18.57** | **23.62** | **22.99** | **22.65** |
| S.E.M. | 3.03 | 3.73 | 1.72 | 2.88 |
| **LACTATE** | **14.53** | **13.57** | **10.17** | **7.16** |
| S.E.M. | 4.25 | 3.12 | 2.66 | 1.98 |
| **MALATE** | **5.24** | **6.81** | **4.98** | **6.04** |
| S.E.M. | 1.54 | 0.33 | 1.00 | 0.89 |
| **PYRUVATE** | **3.36** | **3.26** | **2.93** | **3.02** |
| S.E.M. | 0.65 | 0.56 | 0.45 | 0.64 |
| **SERINE** | **1.10** | **0.94** | **0.75** | **0.99** |
| S.E.M. | 0.28 | 0.08 | 0.08 | 0.12 |
| **SUCCINATE** | **1.24** | **1.24** | **1.26** | **1.66** |
| S.E.M. | 0.20 | 0.12 | 0.06 | 0.25 |
