## Supplemental Table 2 for "^13^C tracing in synaptosomes reveals that SGLT2 inhibition with dapagliflozin prevents metabolic deficits in the 5X-FAD model of Alzheimer’s Disease"

**Supplemental Table 2**: Metabolite abundances in the retrosplenial cortex after two-month Dapagliflozin treatment.

| | BOTH SEXES (nmol/mg protein) | | | |
|---|---|---|---|---|
| | WT | | 5X-FAD Control | |
| | Control | DAPA | Control | DAPA |
| **ALANINE** | **1.65** | **1.66** | **1.72** | **1.88** |
| S.E.M. | 0.10 | 0.14 | 0.14 | 0.18 |
| **ASPARTATE** | **5.76** | **5.11** | **5.39** | **5.79** |
| S.E.M. | 0.40 | 0.41 | 0.38 | 0.64 |
| **CITRATE** | **0.46** | **0.52** | **0.46** | **0.59** |
| S.E.M. | 0.05 | 0.05 | 0.05 | 0.07 |
| **FUMARATE** | **0.30** | **0.32** | **0.28** | **0.35** |
| S.E.M. | 0.03 | 0.03 | 0.02 | 0.04 |
| **GABA** | **3.34** | **3.57** | **3.54** | **4.11** |
| S.E.M. | 0.35 | 0.39 | 0.41 | 0.50 |
| **GLUTAMATE** | **13.69** | **14.62** | **14.53** | **15.56** |
| S.E.M. | 0.73 | 1.42 | 1.27 | 1.46 |
| **LACTATE** | **37.94** | **37.68** | **40.59** | **43.31** |
| S.E.M. | 5.31 | 4.93 | 4.90 | 5.81 |
| **MALATE** | **1.72** | **1.75** | **1.72** | **2.02** |
| S.E.M. | 0.12 | 0.12 | 0.15 | 0.19 |
| **N-acetyl aspa** | **29.17** | **28.21** | **35.12** | **32.40** |
| S.E.M. | 5.56 | 4.76 | 10.46 | 8.07 |
| **PYRUVATE** | **0.28** | **0.27** | **0.27** | **0.27** |
| S.E.M. | 0.03 | 0.02 | 0.02 | 0.03 |
| **SERINE** | **1.53** | **1.50** | **1.62** | **1.81** |
| S.E.M. | 0.12 | 0.11 | 0.15 | 0.18 |
| **SUCCINATE** | **1.24** | **1.32** | **1.44** | **1.43** |
| S.E.M. | 0.10 | 0.13 | 0.15 | 0.16 |

| | FEMALE (nmol/mg protein) | | | |
|---|---|---|---|---|
| | WT | | 5X-FAD Control | |
| | Control | DAPA | Control | DAPA |
| **ALANINE** | **1.59** | **1.80** | **1.65** | **1.72** |
| S.E.M. | 0.21 | 0.22 | 0.25 | 0.25 |
| **ASPARTATE** | **5.53** | **5.26** | **5.36** | **5.39** |
| S.E.M. | 0.79 | 0.69 | 0.42 | 0.86 |
| **CITRATE** | **0.40** | **0.43** | **0.42** | **0.47** |
| S.E.M. | 0.06 | 0.06 | 0.06 | 0.07 |
| **FUMARATE** | **0.28** | **0.31** | **0.26** | **0.30** |
| S.E.M. | 0.05 | 0.03 | 0.04 | 0.06 |
| **GABA** | **1.97** | **2.93** | **2.48** | **2.86** |
| S.E.M. | 0.35 | 0.58 | 0.54 | 0.48 |
| **GLUTAMATE** | **12.92** | **15.91** | **13.43** | **14.49** |
| S.E.M. | 1.39 | 2.53 | 1.63 | 1.97 |
| **LACTATE** | **54.53** | **53.71** | **54.35** | **58.23** |
| S.E.M. | 8.95 | 6.79 | 7.89 | 9.17 |
| **MALATE** | **1.79** | **1.82** | **1.59** | **1.86** |
| S.E.M. | 0.25 | 0.18 | 0.14 | 0.26 |
| **N-acetyl aspa** | **13.06** | **13.67** | **10.35** | **11.00** |
| S.E.M. | 3.07 | 2.77 | 1.24 | 1.85 |
| **PYRUVATE** | **0.28** | **0.29** | **0.28** | **0.26** |
| S.E.M. | 0.04 | 0.02 | 0.03 | 0.03 |
| **SERINE** | **1.96** | **2.12** | **1.94** | **2.09** |
| S.E.M. | 0.25 | 0.33 | 0.22 | 0.28 |
| **SUCCINATE** | **1.24** | **1.33** | **1.20** | **1.39** |
| S.E.M. | 0.21 | 0.15 | 0.14 | 0.15 |

| | MALE (nmol/mg protein) | | | |
|---|---|---|---|---|
| | WT | | 5X-FAD Control | |
| | Control | DAPA | Control | DAPA |
| **ALANINE** | **1.70** | **1.52** | **1.78** | **2.03** |
| S.E.M. | 0.09 | 0.18 | 0.16 | 0.27 |
| **ASPARTATE** | **5.95** | **4.96** | **5.41** | **6.15** |
| S.E.M. | 0.39 | 0.47 | 0.63 | 0.96 |
| **CITRATE** | **0.52** | **0.60** | **0.50** | **0.71** |
| S.E.M. | 0.06 | 0.07 | 0.08 | 0.11 |
| **FUMARATE** | **0.31** | **0.33** | **0.28** | **0.40** |
| S.E.M. | 0.03 | 0.05 | 0.03 | 0.07 |
| **GABA** | **4.44** | **4.20** | **4.40** | **5.24** |
| S.E.M. | 0.20 | 0.46 | 0.47 | 0.68 |
| **GLUTAMATE** | **14.30** | **13.33** | **15.43** | **16.53** |
| S.E.M. | 0.73 | 1.34 | 1.91 | 2.19 |
| **LACTATE** | **24.67** | **21.66** | **29.33** | **29.88** |
| S.E.M. | 1.46 | 2.12 | 3.75 | 4.34 |
| **MALATE** | **1.67** | **1.69** | **1.82** | **2.16** |
| S.E.M. | 0.11 | 0.18 | 0.24 | 0.28 |
| **N-acetyl aspa** | **42.07** | **42.74** | **55.39** | **51.66** |
| S.E.M. | 7.60 | 6.71 | 16.90 | 12.56 |
| **PYRUVATE** | **0.29** | **0.25** | **0.27** | **0.28** |
| S.E.M. | 0.03 | 0.04 | 0.03 | 0.04 |
| **SERINE** | **1.75** | **1.66** | **1.97** | **2.20** |
| S.E.M. | 0.10 | 0.16 | 0.20 | 0.28 |
| **SUCCINATE** | **1.24** | **1.31** | **1.63** | **1.46** |
| S.E.M. | 0.08 | 0.22 | 0.24 | 0.27 |
